# Benchmarking Docking Tools on Experimental and Artificial Intelligence-Predicted Protein Structures

**DOI:** 10.1101/2025.06.03.657620

**Authors:** Paloma Tejera-Nevado, Nathan Junod, Elizabeth Hyunjin Kwon, Lucía Prieto-Santamaría, Alejandro Rodríguez-González

## Abstract

In *silico* analysis provides valuable insights into studying macromolecules, particularly proteins. Protein structure prediction models, like AlphaFold (AF), offer a cost-effective and time-efficient alternative to traditional methods like X-ray crystallography, NMR spectroscopy, and cryo-EM for determining protein structures. These models are increasingly used in protein-ligand interaction studies, a key aspect of drug discovery. Docking and molecular dynamics simulations facilitate this process, and researchers are continuously developing open-access tools for cavity detection and docking to accelerate protein-ligand interaction studies. However, while many of these tools perform well in specific cases, their strengths and weaknesses in analyzing predicted protein structures remain largely unknown. Therefore, it is crucial to compare docking analyses using experimentally determined protein structures and deep learning-based models. In this study, two well-characterized proteins, dopamine D3 receptor with its ligand ETQ and neprilysin with its ligand sacubitrilat, are used to evaluate docking predictions. The docking tools CB-Dock 2 and COACH-D are applied to both X-ray crystallography-derived structures and five different AF-generated models. The objective is to assess the accuracy of these docking approaches and determine whether this strategy can effectively simulate macromolecular behavior in their microenvironment. By doing so, this study aims to generate new insights and contribute to accelerating research in protein-ligand interactions.

## I. INTRODUCTION

Molecular docking is a powerful tool used in drug discovery that allows for in-*silico* elucidation of ligand-protein interactions at an atomic level. There are currently over 100 docking software tools, AutoDock being one of the most widely used [1] [2] [3]. While most current docking tools require user-specification of the ligand binding region, some can identify protein-ligand interactions over the entire protein surface in a process called blind docking [4]. Blind docking can better facilitate exploratory studies of unknown binding modes, and user-friendly web servers such as CB-Dock2 and COACH-D have made these tools even more accessible [5] [6].

Recent development of artificial intelligence (AI)-based tools such as AF 2, have allowed researchers to predict protein structures from sequence data [7]. Using AF-predicted structures in combination with molecular blind docking tools could expand the scope of virtual screening, enabling investigation of target proteins that have not yet been characterized by X-ray crystallography. Because accurate binding site prediction is crucial for the utility of blind docking, several benchmark studies have compared the ability of various docking softwares to reproduce X-ray poses of ligand binding poses [8] [9]. However, the docking performance of ligands onto AF-2 models has been less-extensively characterized and, with current methodologies, demonstrates generally weak performance [10]. There is also a general lack of knowledge regarding which docking tools are best for handling AF-generated structures. Although docking to unmodified AF-2 structures has resulted in less accurate predictions than docking to X-ray crystallography structures [11], the performance slightly improves when docking to holo AF-2 conformations of the protein instead of apo [12].

Since cofactors and small molecules can alter substrate binding interactions and induce conformational changes [13], the inclusion of such molecules into the AF-generated models during docking could improve docking prediction quality. This report investigates this approach by benchmarking the docking of AF-models of two proteins– the dopamine D3 receptor and neprilysin using two web-based blind docking servers– against the crystallized complexes. For this analysis, the performance of two web-based blind docking servers– CB-Dock2 and COACH-D– are compared.

The dopamine D3 receptor (D3R) is a G protein-coupled receptor (GPCR) primarily located in the limbic regions of the brain [14]. It plays a significant role in modulating the mesolimbic dopaminergic pathway, which influences cognition, emotional response, reward mechanisms, and motor control. Dysregulation of D3R signaling has been implicated in several pathological conditions, including Parkinson’s disease, schizophrenia, drug addiction, and depression [15]. Eticlopride (ETQ) is a selective antagonist with high affinity for dopamine D2 and D3 receptors [16]. It has been used heavily in research to investigate receptor-ligand interactions and the specific signaling pathways of dopamine receptors. G protein-coupled receptors (GPCRs) like D3R, are important drug targets because of their heavy involvement in human pathophysiology [17]. Approximately 30% of modern drugs target GPCRs [18]. Advances in computational techniques and AI have enhanced the scope of GPCR drug discovery. Increases in availability and quality of structural data from traditional protein visualization techniques, like X-ray crystallography, have led to the creation of structure-based drug design (SBDB), ligand-based drug design (LBDB), and molecular dynamics (MD) simulations drug discovery workflows. It has also been suggested that, because of the difficulty in capturing the complexity of the intra and extracellular environment *in silico*, the need for human expert consultation or experimental validation using traditional protein identification techniques is required when doing *in silico* drug research [19]. We aim to investigate this claim by docking ligands with PDB files built using X-ray crystallography data and comparing the results to AF predictions made using UniProt sequences from the D3R.

Neprilysin is a type II integral membrane protein and is a zinc-activated endopeptidase involved in various bodily systems, including the cardiovascular, renal, pulmonary, and endocrine systems [20]. Notably, neprilysin breaks down vasodilating peptides and vasoconstrictors angiotensin I and II. Inhibiting neprilysin restores the function of these natriuretic peptides and induces vasodilation, making it a valuable target for congestive heart failure treatment. Sacubitrilat (LBQ657), the active form of sacubitril, is one inhibitor of neprilysin that, when used in combination with valsartan, can reduce risk of cardiovascular death and treat heart failure [21]. Sacubitrilat has been shown to coordinate with the zinc atom when binding to neprilysin [22].

This study aims to compare two docking tools, CB-Dock2 and COACH-D, using data deposited in the PDB database from crystallography studies and ligand information. The comparison focuses on the accuracy of these tools by evaluating their predictions against protein structures generated by AlphaFold, with a specific emphasis on the amino acids predicted to be involved in the cavity and ligand interaction. The first protein analyzed is the dopamine D3 receptor, using experimental data from [14] with the ETQ ligand. Following this, the study examines neprilysin, a protein containing a zinc atom, and its interaction with the ligand sacubitrilat, using experimental data from [22]. By comparing docking outputs with established literature, the research aims to assess the reliability of these prediction tools in identifying relevant protein-ligand interactions, as well as the strength of these prediction tools with the inclusion of biologically relevant small molecules, protein complexes, and ions.

The paper is structured as follows: Section II outlines the tools and resources employed to analyze the cavities and docking. Section III presents the obtained results and discusses them. Lastly, Section IV provides a summary of the study’s conclusions and explores possible directions for analysis in protein-ligand research.

## II. MATERIAL AND METHODS

### A. Protein Sequences and Ligands

The protein sequences were retrieved in PDB format from the PDB databank. The proteins included are the dopamine D3 receptor (UniProt ID: P35462) and neprilysin (UniProt ID: P08473). The corresponding ligands were identified through analysis of the scientific literature, and their structures were retrieved from DrugBank (ETQ: DB15492, Sacubitrilat: DB14127).

AF structures were generated and retrieved in PDB format from the AF server. The query sequences for the dopamine D3 receptor and neprilysin were obtained from UniProt. For the D3R in complex with G proteins and GDP, the G proteins’ (UniProt IDs: P63096, P62873, P63215) sequences were also obtained from UniProt, and a GDP molecule was included in the AF Server query (https://alphafoldserver.com/). For neprilysin, a zinc ion was added to the protein in the AF Server query. All five AF output structures of each query were included in the docking experiments.

To reduce possible interference of extraneous ligands and noise on finding binding pockets, X-ray crystallography-based structures were preprocessed before input to the docking servers. All nonstandard ligands were removed (except for the zinc ion for neprilysin) and the proteins were processed with the DockPrep tool on ChimeraX (v. 1.19) [23] [24].

### B. Protein-Ligand Docking Tools

For cavity and docking analyses, two computational tools, COACH-D and CB-Dock2 [6] [5], were employed. While both tools utilize AutoDock Vina for docking and scoring predictions, they have key differences in their methodologies that could affect docking outcomes. COACH-D relies on sequence-based and template-based predictions, leveraging known protein-ligand complexes to guide docking and cavity identification [6]. In contrast, CB-Dock2 integrates both structure-based cavity detection (blind-dock) and a template-based docking method (fit-dock). CB-Dock2 first detects potential binding cavities from the protein’s structure. If similar templates are found in its database, it proceeds with template-based docking; otherwise, it proceeds with structure-based blind docking without relying on known ligand-binding sites [5]. These methodological differences between COACH-D and CB-Dock2 are critical to consider while interpreting docking predictions and comparing outcomes across studies.

### C. Analysis and Visualization of Docking Results

For protein-ligand visualization, comparison of protein structure, and potential interactions, ChimeraX software (v. 1.9) was used. Different tools, such as Matchmaker, distance measurement, and root mean square deviation (RMSD) [25], were employed to determine similarity between datasets.

To evaluate the success of various docking tools, the accuracy metric was used. This metric calculates the percentage of correctly identified amino acid contact residues predicted when compared to the original amino acids identified using X-ray crystallography by [14] and [22]. The eighteen described amino acids in the Chien et al. study [14] are V86, F106, D110, V111, C114, I183, V189, S192, S193, S196, W342, F345, F346, H349, V350, Y365, T369, Y373. The twelve described amino acids for the interaction of neprilysin with sacubitrilat are R102, R110, F106, N542, F544, H583, E584, H587, E646, W693, G714, and R717 [22].

To examine the interaction between the predicted ligand binding pose and the zinc ion, the closest distance between the ion and one of the ligand’s carboxylate oxygens was measured with the Distances tool in ChimeraX. For predicted complexes containing the zinc ion, this distance was recorded directly. For models without the zinc ion, the complex was first aligned to the corresponding AF protein structure using Matchmaker, and then the distance to the ion was measured.

## III. RESULTS AND DISCUSSION

The docking analyses presented below aim to systematically evaluate how structural complexity and protein assembly of sequences folded by the AF Server AI influence computational predictions from CB-Dock2 and COACH-D. Initially, docking performance of the dopamine D3 receptor is assessed using different structural inputs: single-chain receptor, double-chain receptor, and receptor complexed with G proteins and GDP. This analysis reveals how increasing complexity impacts docking accuracy. Subsequently, the docking of sacubitrilat on neprilysin explores the influence of zinc-ion inclusion on docking outcomes. The combined insights from these analyses underscore the limitations of existing docking tools and highlight the importance incorporating traditionally captured protein structures with AI predictions.

### A. Dopamine D3 receptor as different input models

Docking performance of CB-Dock2 and COACH-D was evaluated against experimental protein structures derived from X-ray crystallography and predictions by AF using sequence data. Docking outcomes differed based on single-chain versus double-chain receptor structures for the dopamine D3 receptor (D3R) and this demonstrated structural input sensitivity.

**Table I** presents the average cavity size and average number of predicted residues compared to [14] for each docking condition. While the cavities created by the cleaned X-ray crystallography PDB files are similar in size, the AF model cavity size increases dramatically as the structures get larger and more complex.

**TABLE 1.** Averages Between Prediction and Receptor Type.

| Prediction Type (PDB) | Number of Proteins | Avg. Cavity Size | Avg. Percent of Correct Residues |
| --- | --- | --- | --- |
| X-ray Ref. | 1 D3R | 907.0 | 58% |
| X-ray Ref. | 2 D3R | 941.0 | 42% |
| AF Model | 1 D3R | 771.4 | 42% |
| AF Model | 2 D3R | 1136.4 | 44% |
| AF Model | 1 D3R, 3 G-proteins | 7378.2 | 32% |

In *silico* analysis of protein and ligand interactions allows for preliminary analyses before conducting further laboratory studies. In our study of the dopamine D3 receptor and its antagonist ETQ, we compared protein structure predictions generated by AF. When using a single-chain input, the results showed that the predictions regarding the amino acids involved in ligand interaction and cavity formation were slightly more accurate compared to using two chains as input (**Table I**). Our results support the hypothesis that integrating traditional experimental methods with AI predictions can improve in silico drug study workflows, as suggested by studies like [19]. The predictive models by themselves can lead to incorrect conclusions, especially when working with complex structures like the D3R in complex with G proteins and GDP (**Table I**). The cleaned PDB files from Chien et al. study [14] performed better than the AF structures due to X-ray crystallography’s better capture of flexible regions and improved binding pocket exposure after ligand removal.

**Table II** compares the results from the cleaned [14] PDB files with the best models produced by the AF server. The X-ray Experiment row shows the experimental data as a control. X-ray Ref. refers to the cleaned X-ray crystallography PDB, and AF Model refers to the best AF server model. Each prediction type is also broken down by the number of proteins included in the docked PDB file. Percent overlap shows the number of residues from the prediction that were shared with the reference structure residues divided by the total number of residues from the Chien et al. study [14]. Accuracy Percentage shows the residues from the prediction that were shared with the Chien et al. study [14] divided by the total number of residues generated by the prediction. This table highlights that the best AF predictions can compete with ligands docked using cleaned X-ray crystallography PDB files.

**TABLE 2.**
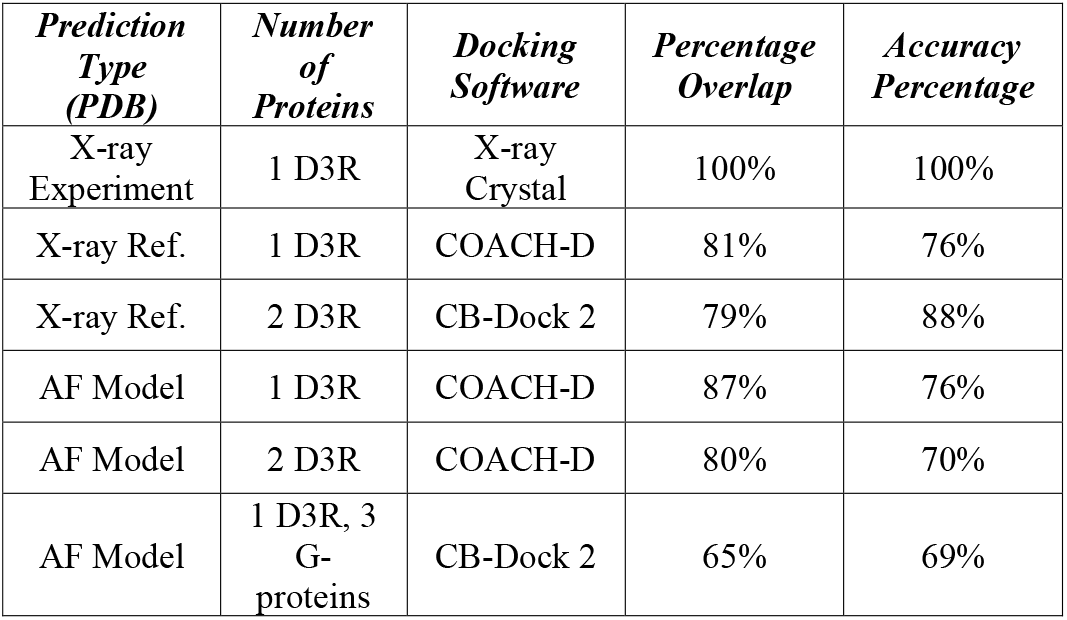
Percentage of Contact Residues Identified By the Best Predictions from each Model Type.

The structural overlap between the top-performing AF-predicted dopamine D3 receptor model docked with COACH-D and the experimentally solved structure [14] is demonstrated in **Fig. 1**. This comparison visualization demonstrates the high degree of structural congruence obtained using AF prediction in the rigid portions of the protein, while also highlighting the flexibility that exists in the loops of the AF prediction.

**Fig. 1.**
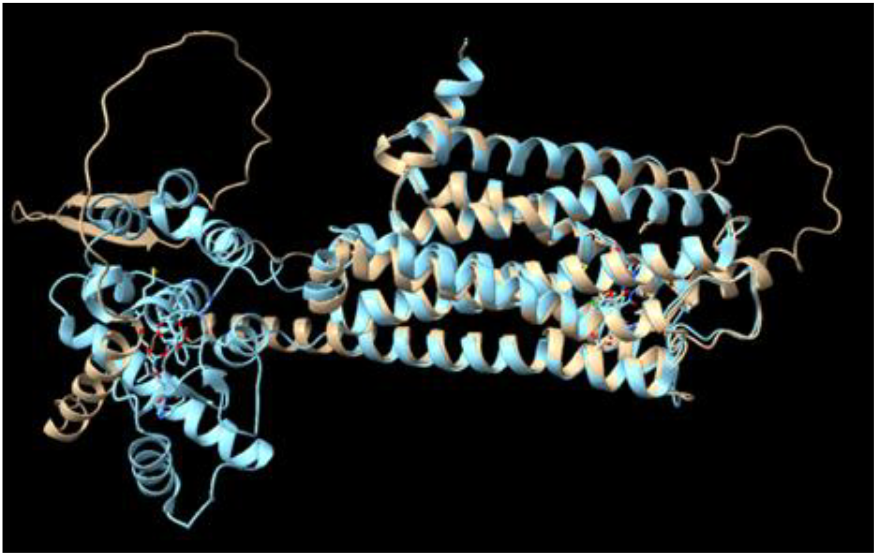
Structural overlap between the top-performing AF-predicted dopamine D3 receptor model docked with COACH-D (beige) and the experimentally solved structure by Chien et al. (2010) (cyan). The sequence alignment score is 1323, and the RMSD between 236 pruned atom pairs is 0.739 (across all 357 pairs: 15.982 Å). These statistics were calculated using the Matchmaker function in Chimera X.

A side-by-side comparison of the best docking predictions using the modified PDB from the reference structure and the original PDB captured by Chien et al. (2010) [14] is shown (**Fig. 2**).

**Fig. 2.**
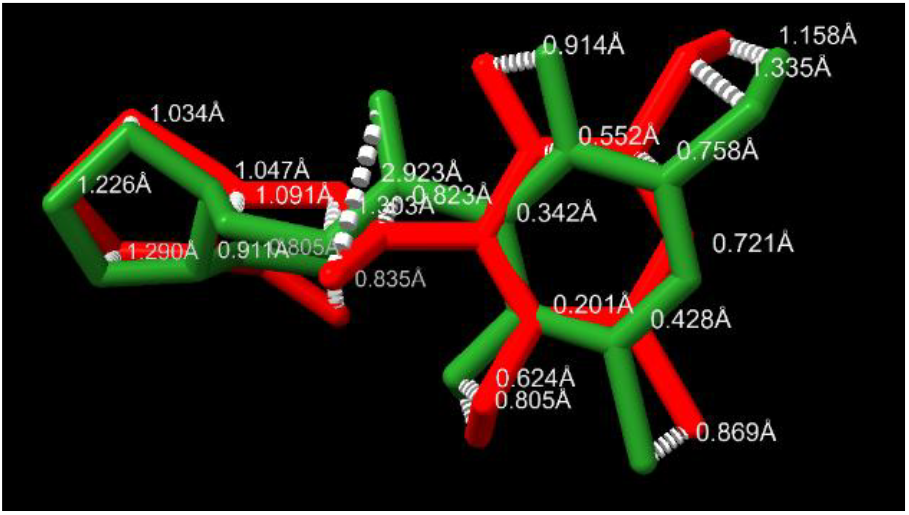
Visualization of the ETQ structure in Chimera X. The dark green ligand represents the original ETQ ligand docked by Chien et al. (2010), while the red ligand represents ETQ docked using the cleaned Chein et al. PDB file with CB-Dock2. Deviations in ligand pose are illustrated by measuring atomic distances in angstroms (Å). The RMSD between atom positions is 6.135 Å, indicating significant differences in ligand positioning between docking methods, particularly in the ketone oxygen near the center of the ETQ ligand.

These findings illustrate the influence of receptor flexibility and the limitations inherent in computational predictions, reinforcing the importance of integrating experimentally validated structures with AI-generated models for accurate and reliable in silico docking analyses (**Fig. 1** and **Fig. 2**). Differences in ligand positioning are observed between the original PDB from Chien et al. (2010) [14] and the cleaned, blind-docked structure (**Fig. 2**). These differences in ligand positioning likely occur due to the flexibility in protein regions, subtle changes in side-chain orientation within the binding pockets, and differences in hydrogen bond interaction calculations. The combined use of traditional and AI approaches is important, as AF models show that proteins, especially GPCRs, are dynamic and flexible in their natural environments. This flexibility means that, while X-ray crystallography provides a good snapshot of the protein at a particular moment, there are many dynamic shifts in protein conformation that we cannot accurately represent or quantify with traditional or AI technologies. Because of this limitation, using AI models like AF to generate multiple models based on varying levels of confidence can help give a more accurate description of the variety of confirmations one may find a particular protein in its environment.

### B. Dopamine D3 receptor in complex with G proteins and GDP

Since it is known that the D3R and G-proteins with GDP form part of a complex, an evaluation of docking cavity prediction was then conducted including the entire complex (**Fig. 3**).

**Fig. 3.**
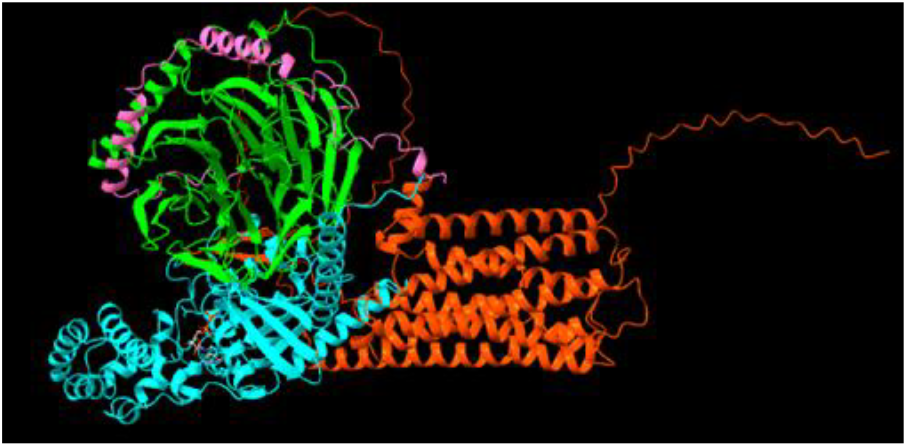
Visualization of the AF prediction of D3R with G-Proteins and GDP in Chimera X. The D3R receptor is represented in red, the Gα subunit in cyan, the Gβ subunit in green, and the Gγ subunit in pink.

Docking predictions involving the D3R complexed with G-proteins and GDP showed notably poorer performance compared to simpler receptor models due to structural complexity and ligand ambiguity. Specifically, the docking tools struggled to identify ETQ-binding residues accurately, as the introduction of the G-proteins (Gα, Gβ, and Gγ subunits) and a GDP molecule significantly altered the receptor’s predicted cavity landscape. The more complex the PDB became, the larger the cavities detected became, causing residue overlap and prediction accuracy to decrease (**Table II**). Consequently, one replicate of the five AF models tested using COACH-D also erroneously focused on GDP rather than ETQ, causing ETQ to bind to the wrong protein chain, reducing the accuracy of overall binding predictions. This finding highlights how overly complex structural inputs, especially involving multiple protein subunits and endogenous ligands such as GDP, can introduce confounding factors that negatively affect the specificity and accuracy of ligand docking simulations. **Table I** also shows the dramatic increase in cavity sizes detected by CB-Dock 2. This increase in cavity size detected is believed to be because the uncertainties in AF prediction models can create large pockets in-between predicted proteins. These pockets are then chosen by the docking software, as they represent a large surface area of available residues. The addition of the GDP binding pocket also created an additional ligand for the software to consider using as a docking candidate. This hybrid approach to docking creates the opportunity for multiple ligand models to be considered during the predictions. For this reason, it seems that keeping files that will be docked as simple as possible will often replicate the best results. The more protein complexes added to the prediction, the more room for error is increased in the system. The dopamine D3 receptor is activated by the G-protein complex (Gα, Gβ, and Gγ) and GDP interaction. Therefore, this complex was also analyzed to determine whether cavity detection shifts with different inputs. The results showed that larger molecular complexes introduced more noise in cavity prediction, placing the potential cavity between the transmembrane domains and increasing the number of ligands that can be used as docking templates.

### C. Protein-Ion-Ligand Analysis

Not all servers could perform docking with zinc included in the protein structure. COACH-D automatically deletes all heteroatoms and ligands from the structure without permitting users to selectively retain them. Consequently, for all five AF structures and the X-ray crystallography query (1DMT), none of the COACH-D outputs included the zinc atom in the output PDB files. In contrast, CB-Dock2 allows selective inclusion of zinc during blind docking. However, successful docking occurred only with AF structures containing the zinc ion, while the X-ray crystallography query (1DMT) failed to dock. As a result, the zinc vs. no-zinc prediction performance could only be compared for the AF-generated models inputted to CB-Dock2.

The docking performance was evaluated using two metrics: first, the number of binding residues shared with the crystallography-determined drug-ligand complex structure [22], as shown in **Table III**; and second, the distances of the ligand carboxylate oxygen to the zinc ion, which were measured as described in the methods section and the results of which are shown in **Table IV**.

**TABLE 3.** Percentage of Predicted Binding Pocket Residues Matching those of the Experimentally Described Complex [22].

| Prediction Type (PDB) | Percentage of correct amino acids (%) |  |  |
| --- | --- | --- | --- |
|  | CB-Dock2 |  | COACH-D |
|  | With Zn<sup>2+</sup> | Without Zn<sup>2+</sup> | Without Zn<sup>2+</sup> |
| AF model 0 | 91.7 | 91.7 | 83.3 |
| AF model 1 | 100 | 83.3 | 83.3 |
| AF model 2 | 100 | 91.7 | 83.3 |
| AF model 3 | 83.3 | 100 | 83.3 |
| AF model 4 | 91.7 | 91.7 | 83.3 |
| X-ray: 1DMT | (*) | 75.0 | 83.3 |

(\*) Option is unavailable

**TABLE 4.** Distance between Zinc ion and closest CArboxylate Oxygen of the Ligand.

| Prediction Type (PDB) | Distance between Zn<sup>2+</sup> and closest carboxylate O (Å) |  |  |
| --- | --- | --- | --- |
|  | CB-Dock2 |  | COACH-D |
|  | With Zn<sup>2+</sup> | Without Zn<sup>2+</sup> | Without Zn<sup>2+</sup> |
| AF model 0 | 5.394 | 5.715 | 5.126 |
| AF model 1 | 4.324 | 1.519 | 4.601 |
| AF model 2 | 2.058 | 1.734 | 3.853 |
| AF model 3 | 1.649 | 5.529 | 2.873 |
| AF model 4 | 2.843 | 4.562 | 6.835 |
| X-ray: 1DMT | (*) | 5.618 | 2.530 |

(\*) Option is unavailable

Some docking yielded higher shared residue counts but larger zinc-ligand distances. For example, AF model 1 (with zinc) docked in CB-Dock2, which covered all reference binding residues but lacked coordination with the zinc ion (**Table III** and **Table IV**). Conversely, other queries resulted in fewer shared residues but closer zinc interactions, such as AF model 3 (with zinc) docked in CB-Dock2 (**Table III and IV**). Since strong performance in one metric did not guarantee strong performance in the other, both interaction distances and shared residues should be considered when docking ligands to proteins with potential metal ion interactions. Despite the five AF neprilysin models exhibiting RMSD values of ≤ 0.188 Å relative to AF model 0, their performance varied across docking servers and zinc conditions. For instance, AF model 2 was the best performing model for both CB-Dock2 conditions (with/without zinc), achieving high shared residue counts and low zinc distances. However, AF model 3 yielded the best COACH-D result without zinc (2.873 Å distance, 10 shared residues). These discrepancies suggest that testing all AF models may be critical for future docking experiments.

To further probe the effect that ligands and ions have on docking predictions, the docking of sacubitrilat on the target neprilysin (with and without the zinc ion) was explored using the X-ray crystallography structure and the AF-generated models. The best generated AF output had an RMSD of 0.282 Å compared to an X-ray crystallography-generated structure of neprilysin (PDB: 1DMT) and retained the zinc atom in the same binding site (**Fig. 4A**). The best predictions for the CB-Dock2 with zinc, CB-Dock2 without zinc, and COACH-D without zinc were obtained from AF model 2, AF model 2, and AF model 3, respectively. Among these, the CB-Dock2 predictions (**Fig. 4B**) had comparable ligand poses (RMSD: 1.159 Å) and slightly outperformed the COACH-D prediction. Thus, for AF structures, CB-Dock2 was a better predictor of the ligand binding pose, regardless of zinc inclusion. Inclusion of zinc did not clearly benefit the docking quality in CB-Dock2 for the AF models. The best predictions in their binding pocket compared to the experimental had an RMSD value of approximately 7.2 Å to the experimental ligand position (**Fig. 4C**). Given the large cavity size (∼1100 Å^3^), this difference could be due to rotational movement within the binding site.

**Fig. 4.**
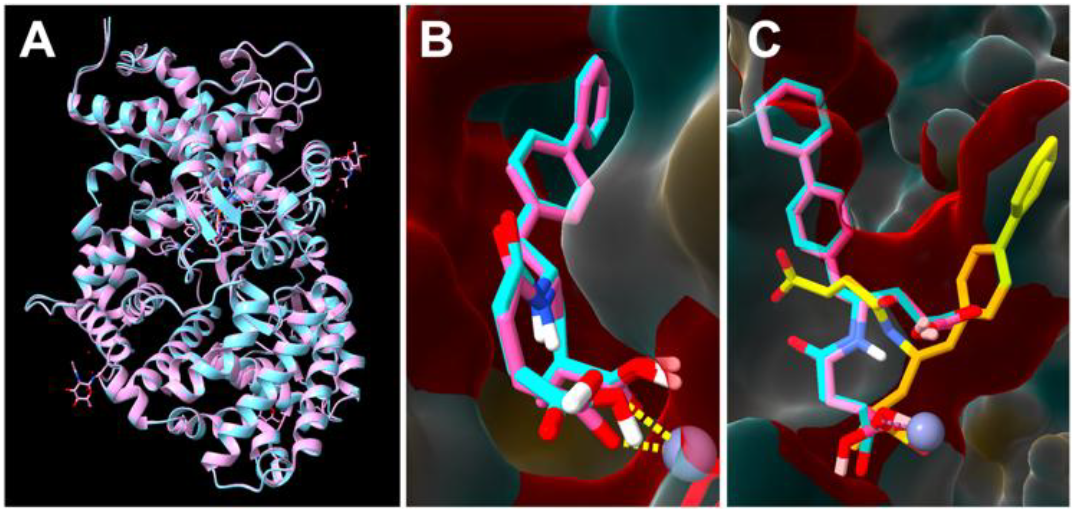
Visualizations of neprilysin CB-Dock2 docking for AF structures including zinc in Chimera X. (A) AF model 0 overlayed with 1DMT via matchmaker. (B) Predicted docking pose from CB-Dock2 on AF 2 (blue – no zinc included, pink – with zinc). The purple sphere at the bottom right is the zinc ion, with dotted yellow lines indicating potential coordination with a carboxylate oxygen from each structure. (C) The same two structures from (B), this time with the experimental binding pose alongside for comparison (yellow). The predicted binding residues that match with the experimental ones are colored in red.

For the cleaned X-ray crystallography structure 1DMT, COACH-D performed better than CB-Dock2, with both a higher number of matching residues and a smaller distance to the zinc atom (**Table III** and **Table IV**). An important research direction would involve systematic investigation into how various cofactors and ions influence docking outcomes when integrated into computational models. Given the observed discrepancies in prediction accuracy when including metal ions such as zinc or sodium, future studies could methodically characterize the impact of different ion types across various protein families. Such systematic characterization would enhance the reliability of computational docking methods, ultimately contributing to more precise and biologically relevant drug discovery pipelines.

## IV. CONCLUSION AND FUTURE WORK

Protein-ligand interaction research is essential in drug understanding biological processes. Traditional methods like X-ray crystallography and NMR spectroscopy provide detailed insights but are time-consuming and expensive. Recent advances in deep learning, particularly neural networks, offer efficient alternatives by predicting binding affinities, docking, and novel binding sites from large datasets. The integration of deep learning with conventional techniques enhances research by quickly screening compounds, which can then be validated experimentally. This combination accelerates drug discovery and opens new possibilities for optimizing therapies and exploring unexplored targets.

Our research supports that a combined approach of using data from docked proteins with data from AI approaches provides the best opportunity to quickly learn new insights about proteins. Building on this, future research should explore the development of drug repurposing pipelines that utilize neural networks trained on extensive datasets combining both experimental and computationally predicted docking interactions. Such pipelines could leverage existing drug data to quickly identify novel therapeutic uses for approved medications, significantly reducing development time and associated costs. Additionally, further enhancement of existing AI protein prediction software is essential. While current tools like AlphaFold have significantly advanced protein structure prediction, increasing the predictive accuracy for complex multi-protein systems, and incorporating dynamic changes induced by ligand binding, remain areas requiring improvement. Future efforts should therefore focus on refining these predictive capabilities, possibly through hybrid models that integrate dynamic molecular simulations with neural network architectures, thus achieving more realistic and functionally relevant structural predictions.

## Acknowledgment

The work is a result of the project “Data-driven drug repositioning applying graph neural networks (3DR-GNN)” that is being developed under grant “PID2021-122659OB-I00” from the Spanish Ministry of Science and Innovation.

